# Gorilla APOBEC3 restricts SIVcpz and influences lentiviral evolution in great ape cross-species transmissions

**DOI:** 10.1101/362079

**Authors:** Yusuke Nakano, Keisuke Yamamoto, Andrew Soper, Ryuichi Kumata, Hirofumi Aso, Naoko Misawa, Yoriyuki Konno, Izumi Kimura, Shumpei Nagaoka, Guillermo Juarez-Fernandez, Jumpei Ito, Terumasa Ikeda, Yoshio Koyanagi, Reuben S Harris, Kei Sato

## Abstract

Restriction factors including APOBEC3 family proteins have the potential prevent cross-species lentivirus transmissions. Such events as well as ensuing pathogenesis require the viral Vif protein to overcome/neutralize/degrade the APOBEC3 enzymes of the new host species. Previous investigations have focused on the molecular interaction between human APOBEC3s and HIV-1 Vif. However, the evolutionary interplay between lentiviruses and great ape (including human, chimpanzee and gorilla) APOBEC3s has not been fully investigated. Here we demonstrate that gorilla APOBEC3G plays a pivotal role in restricting lentiviral transmission from chimpanzee to gorilla. We also reveal that a sole amino acid substitution in Vif is sufficient to overcome the gorilla APOBEC3G-mediated species barrier. Moreover, the antiviral effects of gorilla APOBEC3D and APOBEC3F are considerably weaker than those of human and chimpanzee counterparts, which can result in the skewed evolution of great ape lentiviruses leading to HIV-1.

**Highlights:**

- SIVcpz requires M16E mutation in Vif to counteract gorilla A3G
- Acidic residue at position 16 of Vif is crucial to counteract gorilla A3G
- Gorilla A3D and A3F poorly suppress lentiviral infectivity
- SIVgor and related HIV-1s counteract human A3D and A3F independently of DRMR motif

## Introduction

Lentiviruses were identified in great apes including humans (*Homo sapiens*; HU), central chimpanzees (*Pan troglodytes troglodytes*; CPZ), eastern chimpanzees (*Pan troglodytes schweinfurthii*), and gorillas (*Gorilla gorilla gorilla*; GOR). The lentiviruses isolated from their respective hosts were designated HIV (Barre-Sinoussi et al., 1983), SIVcpzPtt [simian immunodeficiency virus (SIV) in CPZ] (Gao et al., 1999; Keele et al., 2006), SIVcpz*Pts* (SIV in eastern chimpanzee) (Vanden Haesevelde et al., 1996), and SIVgor (SIV in GOR) (Van Heuverswyn et al., 2006). HIV type 1 (HIV-1) is classified into four groups, M (major), N (non-M-non-O), O (outlier) and P (reviewed in (Sharp and Hahn, 2011). Molecular phylogenetic analyses indicate that that HIV-1M and HIV-1N originated from SIVcpz*Ptt* (Keele et al., 2006), while HIV-1O and HIV-1P are derived from SIVgor (D’arc et al., 2015). These insights indicate that HIV-1s emerged by cross-species viral transmission from CPZ and GOR, respectively.

To potently control cross-species lentiviral transmission, several cellular restriction factors such as TRIM5, tetherin, SAMHD1 and APOBEC3 (A3) proteins were identified [reviewed in (Doyle et al., 2015; Kluge et al., 2015)]. One of the well-studied restriction factors that potently restricts cross-species transmission of great ape lentiviruses is HU tetherin. To overcome the HU tetherin-mediated antiviral effect, the viral protein U (Vpu) of HIV-1M, a pandemic virus group, down-modulates and antagonizes HU tetherin (Neil et al., 2008; Van Damme et al., 2008). Since the Vpu protein of SIVcpz*Ptt*, the ancestral virus of HIV-1M (Keele et al., 2006), is incapable of counteracting HU tetherin (Sauter et al., 2009), these observations imply that HU tetherin functions as barrier restricting cross-species lentiviral transmission from CPZ to HU, and that acquiring anti-HU tetherin activity is essential for the successful cross-species jump [reviewed in (Kirchhoff, 2010)].

Another group of well-understood restriction factors is the A3 DNA deaminase family. Most great apes encode seven A3 proteins [reviewed in (Nakano et al., 2017a)]. At least three, A3D, A3F, and A3G, are packaged into nascent viral particles and suppress viral infectivity through inserting G-to-A mutations in the viral genome [reviewed in (Harris and Dudley, 2015)]. To counteract A3-mediated restriction action, viral infectivity factor (Vif), an accessory protein of lentiviruses, recruits cellular E3 ubiquitin ligase complex and degrades the host A3 proteins via a ubiquitin-proteasome-dependent pathway [reviewed in (Harris and Dudley, 2015)]. Because Vif-mediated counteraction of antiviral A3 is largely species-specific, it is suggested that lentiviral *vif* and mammalian *A3* genes have co-evolve [reviewed in (Nakano et al., 2017a)]. For instance, it has been reported that Old world monkey A3G proteins contribute to restrict lentiviral transmission among these species (Compton et al., 2014; Compton and Emerman, 2013).

SIVgor was first discovered from the fecal samples of wild GORs, which were obtained in remote forest regions in Cameroon in 2007 (Van Heuverswyn et al., 2006). A following study revealed that SIVgor is phylogenetically related to HIV-1 groups O and P (D’arc et al., 2015), suggesting that SIVgor is the ancestral virus of these HIV-1s in the human population. Moreover, a phylogenetic analysis deduced that SIVgor emerged from the leap of SIVcpz*Ptt* from CPZ to GOR (Takehisa et al., 2009).

The molecular interactions between HIV-1 Vif and HU A3 proteins are well established [reviewed in (Harris and Dudley, 2015)]. Also, the functional and evolutionary relationships between SIVcpz*Ptt* Vif and HU A3 proteins have been investigated (Letko et al., 2013; Sato et al., 2018; Zhang et al., 2017). However, the evolutionary episodes of great ape lentiviruses through GOR, namely, the emergence of (i) SIVgor from SIVcpz*Ptt*; and (ii) HIV-1OP from SIVgor, have not been fully investigated. Moreover, the functional and evolutionary association of great ape lentiviral Vif with their host A3 proteins remains unclear. In this study, we focus on the antiviral effects of great ape A3 proteins including GOR A3s and their antagonistic mechanisms by great ape lentiviral Vif proteins. To the best of our knowledge, this is the first report suggesting that a great ape A3 protein potently restricts the cross-species leap of great ape lentiviruses and illustrates the evolutionary scenario of great ape lentiviruses via the interaction of great ape A3 proteins.

## Results

### GOR A3G restricts SIVcpz*Ptt* infection

Figure 1A illustrates a phylogenetic tree of *vif* genes of great ape lentiviruses. This phylogenetic tree indicates that HIV-1M and HIV-1N form clusters with SIVcpz*Ptt*, while HIV-1O and HIV-1P are together with SIVgor. Consistent with previous reports, it is suggested that HIV-1M and HIV-1N originated from SIVcpz*Ptt* (Keele et al., 2006), while HIV-1O and HIV-1P are derived from SIVgor (D’arc et al., 2015) (Figure 1B).

**Figure 1.**
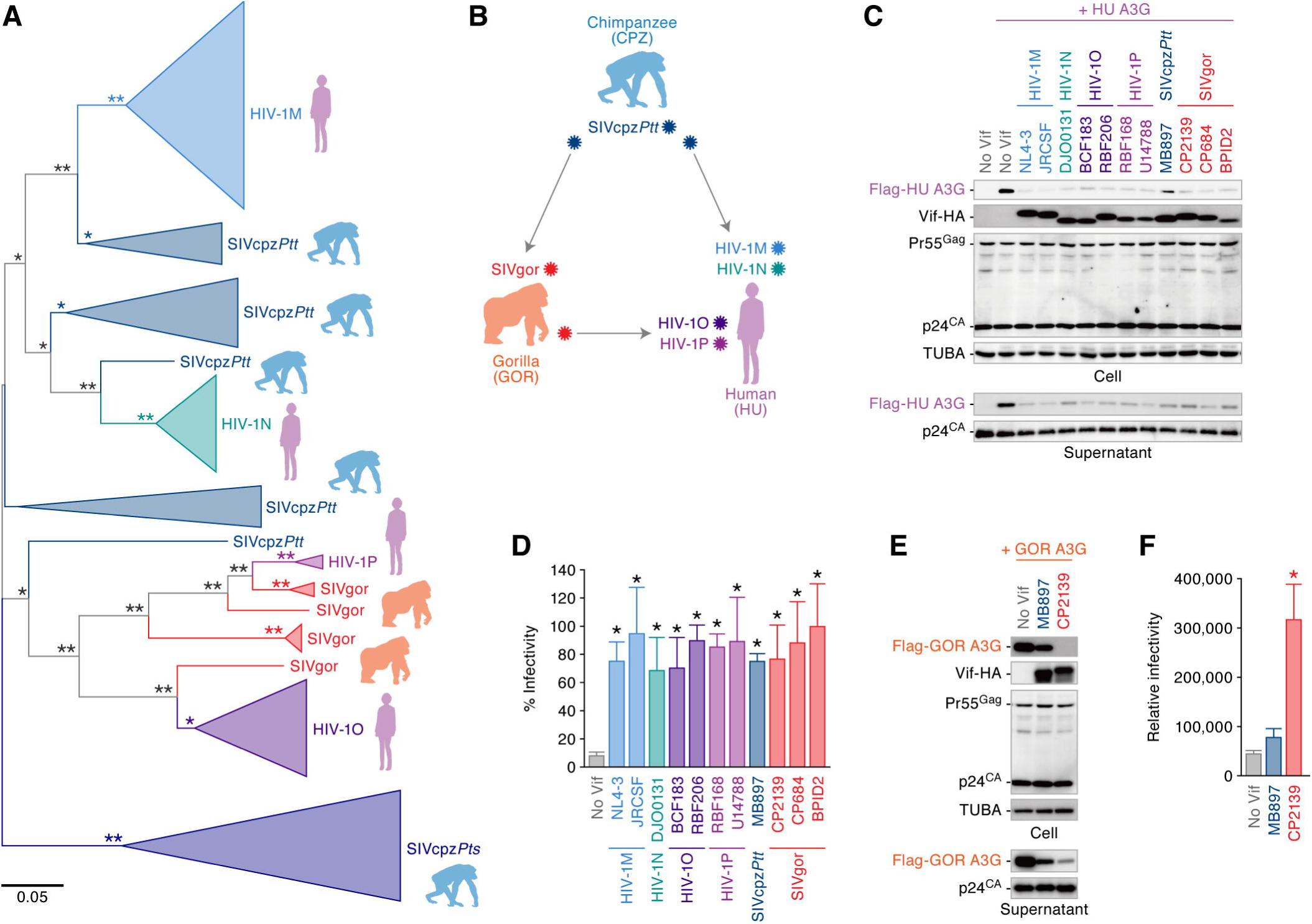
Cross-species lentiviral transmission in great apes. (A) Phylogenetic tree of the *vif* gene of great ape lentiviruses. The *vif* sequences were extracted from Los Alamos HIV sequence database (https://www.hiv.lanl.gov/components/sequence/HIV/search/search.html) and the phylogenetic tree was constructed as described in **STAR^*^METHODS**. The bootstrap values are indicated as follows: ^*^, >50%; ^**^, >80%. Scale bar indicates 0.05 nucleotide substitutions per site. An uncollapsed tree is shown in **Figure S1A**, and the accession numbers of viral sequences are summarized in **Table S1**. (B) Scheme of cross-species lentiviral transmission in great apes. (C and D) Anti-HU A3G activity of primate lentiviral Vif. HEK293T cells were co-transfected with pNL4-3Δ*vif* and the expression plasmids for HU A3G and indicated Vif. Cells and supernatants were harvested at two days post-transfection and were used for Western blotting (C) and TZM-bl assay (D). (E and F) Counteracting ability of SIVcpz and SIVgor Vif against GOR A3G. HEK293T cells were co-transfected with pNL4-3Δ*vif* and the expression plasmids for GOR A3G and the Vif of indicated viral strain. Cells and supernatants were harvested at two days post-transfection and were used for Western blotting (E) and TZM-bl assay (F). For Western blotting, the input of cell lysate was standardized to TUBA, and representative results are shown. For TZM-bl assay, the percentage of the value without HU A3G (D) and the raw value (relative infectivity) determined by TZM-bl assay (F) are shown. The mean values of nine independent experiments ± SEM are shown, and statistically significant differences (*P* < 0.05) versus “no Vif” are shown by asterisks. See also **Figure S1** and **Table S1**.

To address the possibility that great ape A3G can be a factor restricting cross-species transmission of great ape lentiviruses, we set out to analyze the antiviral activity of great ape A3G. We co-transfected the expression plasmids of great ape (HU, CPZ and GOR) A3G with an infectious molecular clone (IMC) of *vif*-deleted HIV-1. All great ape A3Gs exhibited comparably strong and dose-dependent antiviral effects (**Figure S1B and S1C**).

To assess the ability of lentiviral Vif to counteract host A3G, the expression plasmid for HU A3G was co-transfected together with the Vif expression plasmids and an IMC of *vif*-deleted HIV-1. As shown in Figure 1C, all lentiviral Vifs including HIV-1MNOP, SIVcpz*Ptt* and SIVgor degraded HU A3G and impaired the incorporation of HU A3G into the released virions. Also, in the presence of HU A3G, the viral infectivity was rescued by all lentiviral Vifs tested in this study (Figure 1D). These findings suggest that HU A3G does not restrict cross-species transmission of SIVs from CPZ and GOR to HU. In sharp contrast, we found that GOR A3G was antagonized by SIVgor Vif but not by SIVcpz*Ptt* Vif (Figures 1E and 1F). These findings strongly suggested that the antiviral activity of GOR A3G had to be overcome for cross-species transmission of SIVcpz*Ptt* from CPZ to GOR.

### M16E mutation confers anti-GOR A3G ability on SIVcpz*Ptt* Vif

To investigate how SIVgor acquired the ability to counteract GOR A3G, we next performed gain-of-function experiments based on SIVcpz*Ptt* Vif (strain MB897). We first prepared four chimeric Vif mutants of SIVcpz*Ptt* MB897 and SIVgor CP2139 (chimeras A-D; Figure 2A) and evaluated their ability to counteract GOR A3G-mediated antiviral action using the cell-based single-round infection assays. As shown in Figure 2B, SIVgor CP2139 Vif as well as chimera A Vif overcame the GOR A3G-mediated antiviral effect, suggesting that the N-terminal domain of SIVgor is responsible for the counteraction of GOR A3G. We then prepared five additional mutants (chimeras A1-A5) and performed cell-based co-transfection experiments. We found that only chimera A1 is able to degrade GOR A3G (Figure 2C, **top**) and therefore rescues the infectivity of released viruses (Figure 2C, **bottom**). Only three amino acid differences occur in this region (**Figure S2A**). Individual mutants were analyzed and only M16E, not the K6Q or D14P, conferred the ability to counteract GOR A3G (Figure 2D). To analyze the effect of M16E mutation on viral spread, we constructed the mutant IMCs of SIVcpz*Ptt* MB897 and prepared the viral supernatant. The infectivity of the M16E and E2X (*vif*-deleted) variants was comparable to wild-type (WT) virus in the absence of A3s (**Figure S2B**). We inoculated these viruses into the HU A3-null SupT11-CCR5 cells and the SupT11-CCR5 cells stably expressing GOR A3G (SupT11-CCR5-GOR A3G) (**Figure S2C**). In parental SupT11-CCR5 cells, these three viruses replicated similarly (Figure 2E, **left**). On the other hand, in SupT11-CCR5-GOR A3G cells, the growth kinetics of the M16E mutant was significantly higher than those of WT and the E2X mutant (Figure 2E, **right**). Moreover, with regard to the area under the curve indicating the amount of cumulative viruses released in the culture supernatant, this value of the M16E mutant was significantly higher than those of WT and the E2X mutant in SupT11-CCR5-GOR A3G cells (Figure 2F). These findings show that Vif position 16 is an important determinant of GOR A3G counteraction.

**Figure 2.**
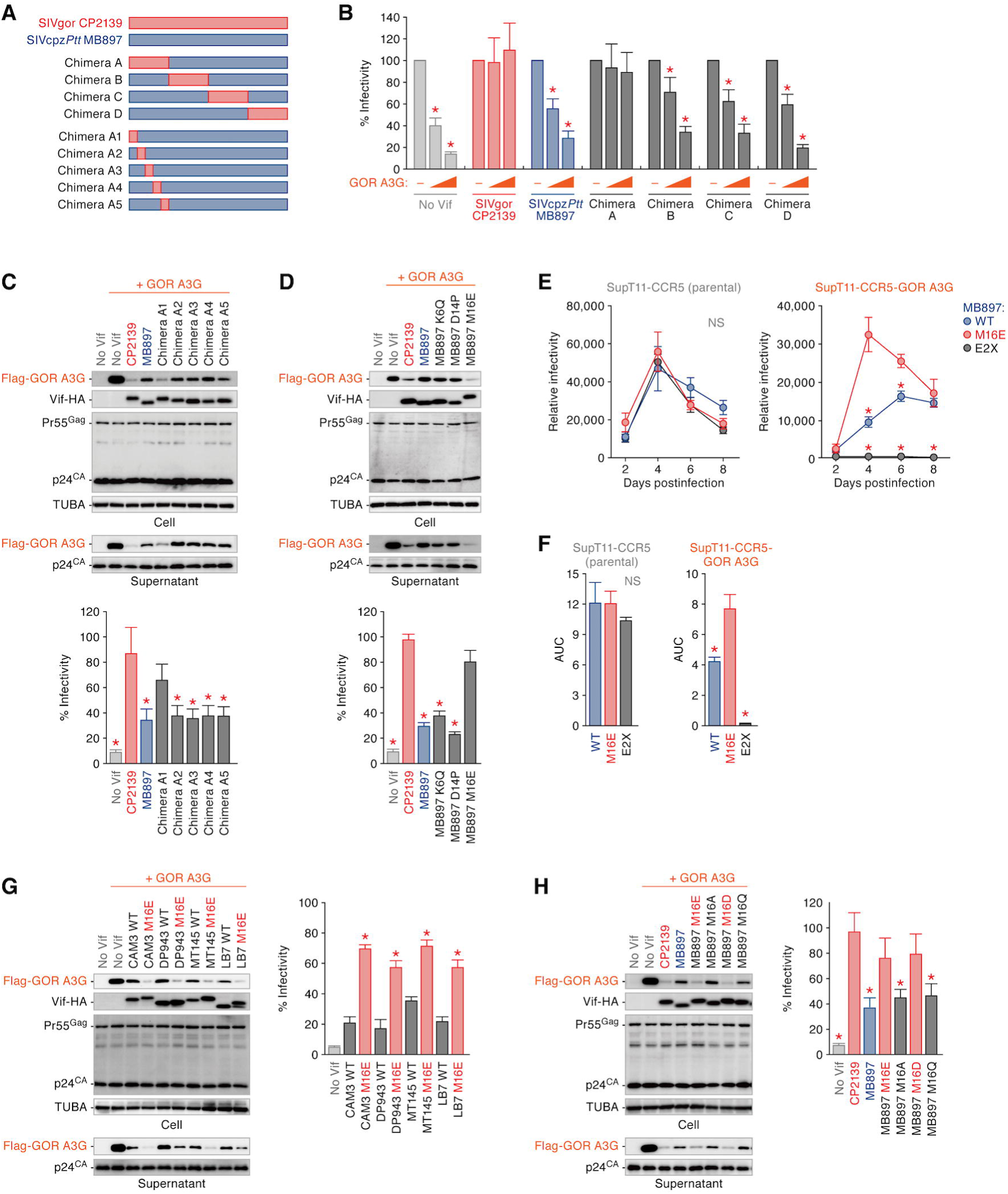
Gain-of-function of SIVcpz*Ptt* Vif to counteract GOR A3G. (A) Scheme of SIVcpz*Ptt* MB897 Vif derivatives used in this study. The amino acid sequences of respective mutants are shown in **Figure S2A**. (B-D) Determination of the amino acid residue of SIV Vif that is responsible to counteract GOR A3G. HEK293T cells were co-transfected with pNL4-3Δ*vif* and the expression plasmids for GOR A3G (10 or 50 ng; the plasmid amount was normalized by empty vector) and the indicated Vif derivatives. Cells and supernatants were harvested at two days post-transfection and were used for Western blotting and TZM-bl assay. (E and F) Multi-round replication assay. The infectious viruses of SIVcpz*Ptt* MB897 WT, M16E and E2X (*vif* deleted) derivatives were prepared as described in **STAR^*^METHODS** and were inoculated into parental SupT11-CCR5 cells or the SupT11-CCR5 stably expressing GOR A3G (see also **Figure S2B**) at MOI 0.1. (E) Culture supernatant was harvested per two days and the amount of infectious viruses was measured by using TZM-bl cells. (F) Area under the curve (AUC) of respective virus infection culture is shown. (G) Importance of M16E substitution in other SIVcpz*Ptt* strains to counteract GOR A3G. (H) Importance of acidic residue at position 16 of SIVcpz*Ptt* to counteract GOR A3G. In (G) and (H), HEK293T cells were co-transfected with pNL4-3Δ*vif* and the expression plasmids for GOR A3G and the indicated Vif strains and derivatives. Cells and supernatants were harvested at two days post-transfection and were used for Western blotting (left) and TZM-bl assay (right). For Western blotting, the input of cell lysate was standardized to TUBA, and representative results are shown. For TZM-bl assay, the percentage of the value without GOR A3G is shown. The mean values of nine independent experiments ± SEM are shown. Statistically significant differences (*P* < 0.05) versus “CP2139” (B-D, G and H) or “M16E” (E and F) are shown by red asterisks. NS, no statistical significance. See also **Figure S2**.

As shown in **Figure S2D**, M16 in Vif is conserved among 16 different SIVcpz*Ptt* strains. To ask whether the substitution of methionine to glutamic acid broadly contributes to the gain-of-function of SIVcpz*Ptt* Vif to counteract GOR A3G, we constructed M16E mutants of additional SIVcpz*Ptt* strains, CAM3, DP943, MT145 and LB7. Although the WT Vifs of these SIVcpz*Ptt* strains were unable to counteract GOR A3G, all M16E mutants of SIVcpz*Ptt* Vif tested in this experiment acquired the ability to counteract GOR A3G (Figure 2G). These findings suggest that the acquisition of anti-GOR A3G ability by the M16E substitution is not specific for SIVcpz*Ptt* strain MB897 but shared among SIVcpz*Ptt* strains.

We next assessed the side-chain properties of position 16 of SIVcpz*Ptt* Vif to counteract GOR A3G. To address this, we constructed three additional mutants at position 16. In addition to the M16E mutant, the M16D mutant of MB897 Vif counteracted the GOR A3G-mediated antiviral effect, while the M16A and M16Q mutants did not (Figure 2H). Taken together, these results suggested that acquiring an acidic residue at position 16 is sufficient to counteract GOR A3G-mediated antiviral effect.

### GOR A3D and A3F poorly exhibit antiviral effect

It is known that the DRMR motif, residues positioned between 14-17 in Vif, are crucial to counteract HU A3D and A3F (Russell et al., 2009; Sato et al., 2014). Since M16E is located in this motif, we next investigated the association of GOR A3D and A3F with great ape lentiviruses. We co-transfected the expression plasmids of great ape A3D or A3F with an IMC of *vif*-deleted HIV-1. Interestingly, the expression levels of GOR A3D (Figure 3A, **top**) and A3F (Figure 3C, **top**) were clearly lower than those of HU and CPZ counterparts. Also, the amounts of GOR A3D (Figure 3A, **bottom**) and A3F (Figure 3C, **bottom**) in the released viral particles as well as the antiviral effects of GOR A3D (Figure 3B) and A3F (Figure 3D) were lower than those of HU and CPZ counterparts. These data suggest that GOR A3D and A3F are poorly expressed and faintly exhibit antiviral effect when compared to HU and CPZ counterparts.

**Figure 3.**
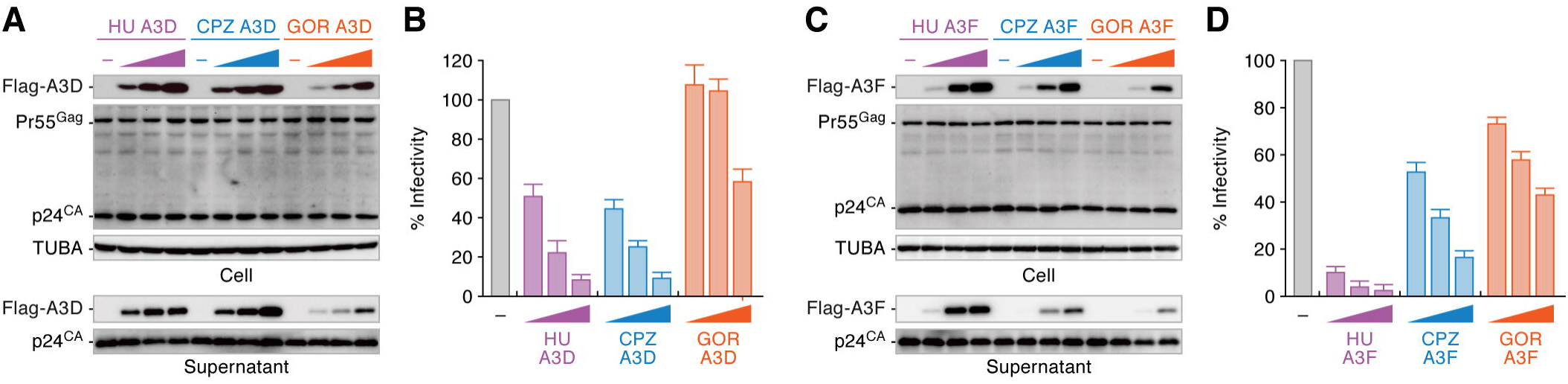
Poor antiviral activity of GOR A3D and A3F. Antiviral activity of great ape A3D (A and B) and A3F (C and D). HEK293T cells were co-transfected with pNL4-3Δ*vif* and the different amounts of expression plasmids for great ape A3D and A3F (0, 50, 100, and 200 ng; the plasmid amount was normalized by empty vector). Cells and supernatants were harvested at two days post-transfection and were used for Western blotting (A and C) and TZM-bl assay (B and D). For Western blotting, the input of cell lysate was standardized to TUBA, and representative results are shown. For TZM-bl assay, the infectivity value without A3 was set to 100%.

### Vif DRMR motif is distortedly evolved in GOR

We next assessed the conservation of the DRMR motif in great ape lentiviral Vif. As shown in Figure 4A, this motif is highly conserved in HIV-1M, HIV-1N, SIVcpz*Ptt* and SIVcpz*Pts*, another group of SIVcpz in a subspecies of chimpanzee (eastern chimpanzee; *Pan troglodytes schweinfurthii*). In contrast, this domain is altered in HIV-1O, HIV-1P and SIVgor (Figure 4A). Moreover, the molecular phylogenetic analysis on the Vif DRMR motif indicated that this motif in HIV-1O, HIV-1P and SIVgor forms separated clusters to HIV-1M, HIV-1N, SIVcpz*Ptt* and SIVcpz*Pts* with higher bootstrap values (**Figure S3A**). Since the YRHHY motif positioned between 40-44 in Vif, which is responsible for A3G counteraction (Russell et al., 2009; Sato et al., 2014), is highly conserved in all great ape lentiviruses (Figure 4A), these findings suggest that the Vif DRMR motif may be less functionally constrained.

**Figure 4.**
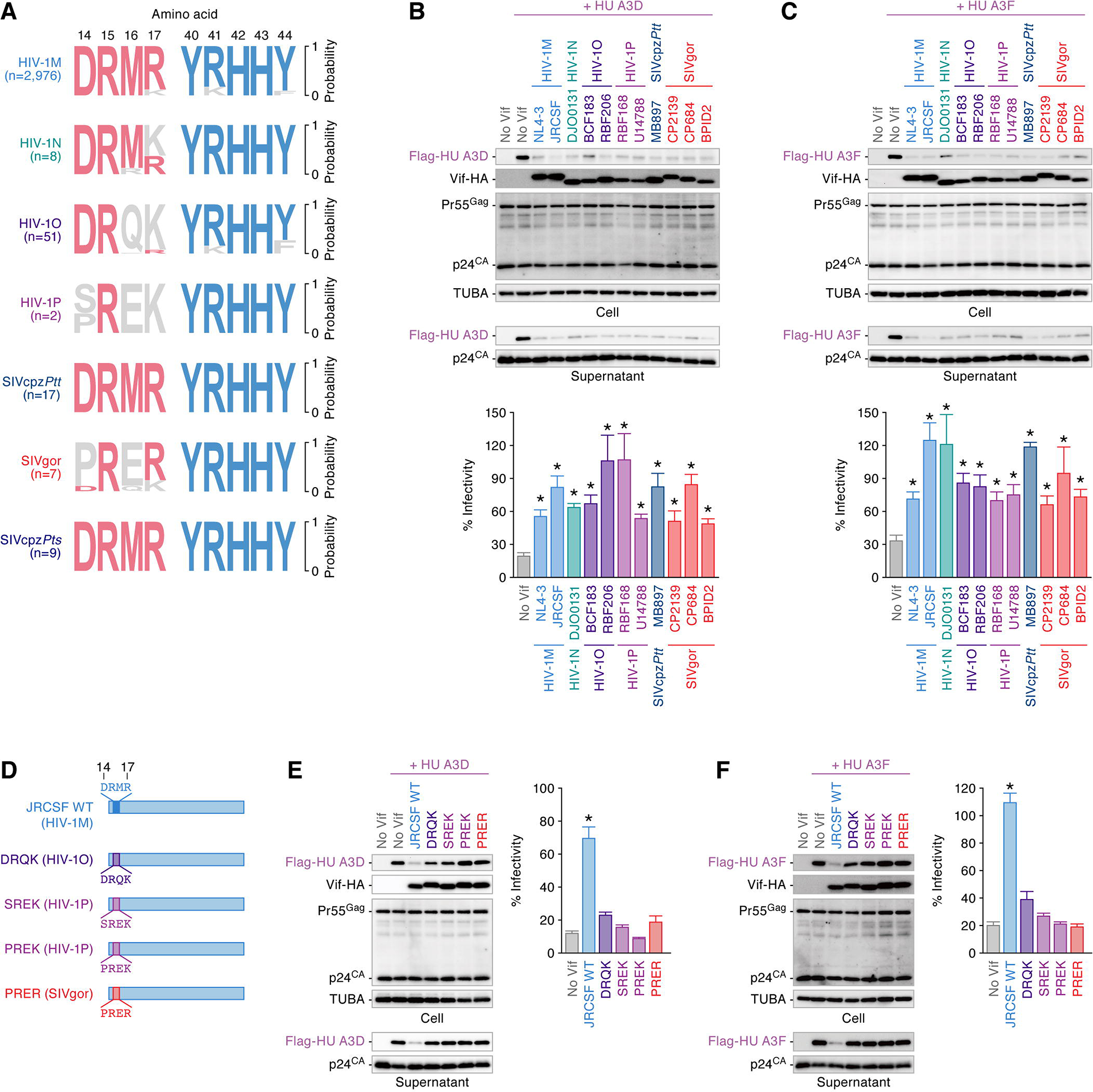
Degradation of antiviral human A3D and A3F by HIV-1OP and SIVgor Vif independently of DRMR motif. (A) Relaxed selection in DRMR motif of SIVgor and related HIV-1 groups. Logo plot of DRMR and YRHHY motifs in HIV-1MNOP, SIVcpz and SIVgor are shown. The number in parenthesis (n) indicates the number of viral sequences used in this analysis. (B and C) Activity of primate lentiviral Vif against HU A3D and A3F. HEK293T cells were co-transfected with pNL4-3Δ*vif* and the expression plasmids for HU A3 and indicated Vif. Cells and supernatants were harvested at two days post-transfection and were used for Western blotting (top) and TZM-bl assay (bottom). (D) Scheme of HIV-1M JRCSF Vif derivatives used in this study. (E and F) Vif’s activity against HU A3D and A3F. HEK293T cells were co-transfected with pNL4-3Δ*vif* and the expression plasmids for HU A3D (E) or A3F (F) and indicated Vif. Cells and supernatants were harvested at two days post-transfection and were used for Western blotting (left) and TZM-bl assay (right). For Western blotting, the input of cell lysate was standardized to TUBA, and representative results are shown. For TZM-bl assay, the percentage of the value without HU A3 is shown. The mean values of six independent experiments ± SEM are shown, and statistically significant differences (*P* < 0.05) versus “no Vif” are shown by asterisks. See also **Figure S3**.

The loss of the DRMR motif in the Vifs of HIV-1O, HIV-1P and SIVgor raised the possibility that these Vifs are unable to counteract A3D and A3F from HU and GOR. To address this issue, the expression plasmids for HU A3D or A3F were co-transfected together with the Vif expression plasmids and an IMC of *vif*-deleted HIV-1. As shown in Figures 4B and 4C, it was surprising that the all Vifs tested including those from HIV-1O, HIV-1P and SIVgor counteracted both HU A3D and A3F. These Vifs also counteracted GOR A3D and A3F (**Figure S3B and S3C**).

These findings (Figures 4A–4C) suggested that Vif amino acids 14-17 in HIV-1O, HIV-1P and SIVgor may be dispensable for counteracting HU A3D and A3F. To address this possibility, we constructed chimeric mutants of HIV-1M strain JRCSF Vif that possess the amino acid residues of HIV-1O, HIV-1P and SIVgor at the corresponding positions: DRQK (from HIV-1O), SREK (from HIV-1P), PREK (from HIV-1P) and PRER (from SIVgor) (Figure 4D). Although the Vifs of HIV-1O, HIV-1P and SIVgor successfully counteracted HU A3D and A3F (Figures 4B and 4C), the JRCSF Vif mutants possessing the amino acid residues of HIV-1O (DRQK), HIV-1P (SREK and PREK) and SIVgor (PRER) instead of DRMR were unable to counteract HU A3D (Figure 4E) and A3F (Figure 4F). Taken together, these findings suggest that HIV-1O, HIV-1P and SIVgor overcome the antiviral effects mediated by A3D and A3F of HU and GOR independently of the Vif DRMR motif.

## Discussion

In the present study, we demonstrated that HU A3D, A3F, and A3G are counteracted by all great ape lentiviral Vifs tested in this study. Our results are consistent with previous observations that SIVcpz (Bibollet-Ruche et al., 2012) and SIVgor (Takehisa et al., 2009) are able to replicate in *in vitro* human CD4^+^ T-cell culture. Moreover, SIVcpz efficiently expands in a hematopoietic stem cell-transplanted humanized mouse model (Sato et al., 2018; Yamada et al., 2018). Taken together with our findings, these observations suggest that HU A3 proteins may not have been a barrier to SIV transmission from CPZ and GOR to HU. In contrast, SIVcpz*Ptt* Vif was incapable of antagonizing the GOR A3G-mediated antiviral effect. Here we demonstrated that an amino acid substitution at position 16 of SIVcpz*Ptt* Vif is sufficient to acquire the ability to counteract GOR A3G. Our findings suggest that GOR A3G restriction activity had to be overcome for transmission from of SIVcpz*Ptt* from CPZ to GOR. To the best of our knowledge, this is the first report providing evidence that a great ape A3 protein plays a crucial role in restricting the leap of great ape lentivirus between different host species and how exactly primate lentiviruses overcame the hurdle mediated by host antiviral A3 protein. Furthermore, we revealed that the antiviral effects of GOR A3D and A3F are relatively weaker than those of HU and CPZ counterparts and that the Vif DRMR motif, which is important to counteract HU A3D and A3F (Russell et al., 2009; Sato et al., 2014), has been lost in SIVgor and related HIV-1s (groups O and P). These findings suggest that the evolutionary process of great ape lentiviruses leading to HIV-1 in HU was uniquely skewed via SIV infection in GOR.

With regard to the functional interaction between GOR A3G and the Vif proteins of SIVcpz*Ptt* and SIVgor, a previous paper revealed that the amino acid positioned at 129 of GOR A3G determines the sensitivity to SIVgor Vif-mediated degradation (D’arc et al., 2015). Interestingly, only GOR A3G possesses glutamine at position 129 while the A3G of other primates including HU and CPZ possesses proline at that position (Letko et al., 2015; Letko et al., 2013). However, because of the low sequence similarity on the *vif* genes of SIVcpz*Ptt* and SIVgor (69.6% nucleotide sequence similarity between SIVcpz*Ptt* strain MB897 and SIVgor strain CP2139, both of which were used in this study for chimeric Vif preparation [Figure 2A]; p-distance between SIVcpz**Ptt** *vif* and SIVgor *vif* = 0.373 ± 0.013), the responsible residue(s) in Vif determining the ability to counteract GOR A3G were not revealed. Here we demonstrated that the ability of SIVcpz*Ptt* Vif to counteract GOR A3G is acquired by a single amino acid substitution at the position 16. Interestingly, with regard to the structural interaction between Vif and A3G, Letko et al. have recently provided a co-structure model of Vif and A3G suggesting that the amino acid residues located between 14-17 of Vif structurally interacts with the residues located between 125-130 of A3G and that the electric charge of the surface of respective domains on each protein associates with Vif-A3G interaction (Letko et al., 2015). Taken together with our finding that the acquisition of an acidic residue (glutamic acid or aspartic acid) at position 16 of SIVcpz**Ptt** Vif confers the ability to counteract GOR A3G, the electrostatic interaction between these domains may be crucial for the functional interaction between Vif and GOR A3G.

Surprisingly, the amino acid residue responsible for the counteraction of GOR A3G was located in the Vif DRMR motif, which is required for degradation of HU A3D and A3F (Russell et al., 2009; Sato et al., 2014). Additionally, the Vif DRMR motif was less conserved in SIVgor and related HIV-1 (groups O and P), while this motif was highly conserved in SIVcpz*Ptt* and related HIV-1 (groups M and N). Moreover, the phylogenetic tree of the Vif DRMR motif revealed that SIVgor forms a cluster with HIV-1 groups O and P. These findings indicated that the Vif DRMR motif, in particular, has been lost when GOR was the host species. In this regard, we demonstrated that the antiviral activity of GOR A3D and A3F is relatively weaker than that of HU and CPZ counterparts. Taken together with these findings, the skewed evolution of the Vif DRMR motif in GOR might be permitted due to the selection pressure mediated by A3D and A3F being relaxed in GOR. Furthermore, although the *vif* nucleotide sequence similarity between SIVcpz*Ptt* and SIVcpz*Pts*, which is classified as the outgroup of HIV-1, SIVcpz*Ptt* and SIVgor, was relatively low (p-distance between SIVcpz*Ptt vif* and SIVcpz*Pts vif* = 0.347 ± 0.016), the DRMR motif is highly conserved in SIVcpz*Pts* and the phylogenetic tree of the Vif DRMR motif indicated that SIVcpz*Pts* forms a cluster with SIVcpz*Ptt* and related HIV-1s. These observations further suggest that the loss of the Vif DRMR motif may be specific to SIVgor and related viruses.

Although SIVgor and related HIV-1s lost the Vif DRMR motif in GOR, these Vifs showed the ability to counteract A3D and A3F from GOR and HU. These findings suggest that the loss of the DRMR motif resulted in the acquisition of novel domain(s) to counteract A3D and A3F and to maintain Vif’s integrity. In addition to the distorted evolution in the Vif DRMR motif, previous studies suggested that other viral antagonists, Vpu and Nef, have uniquely evolved in GOR: based on the findings that HIV-1M Vpu but not SIVcpz*Ptt* Vpu counteracts HU tetherin (Sauter et al., 2009), it is implied that gain of anti-HU tetherin activity by Vpu is important for cross-species lentiviral transmission from CPZ to HU [reviewed in (Kirchhoff, 2010)]. However, Kluge et al. revealed that HIV-1O Nef but not Vpu has gained the ability to counteract HU tetherin (Kluge et al., 2014). Since HIV-1O emerged from SIVgor (D’arc et al., 2015), this is another example of the unique evolution of lentiviral genes in GOR. Therefore, the unique lentiviral evolution in GOR including the distortion of the Vif DRMR motif may contribute to the relatively lower prevalence of HIV-1 groups O and P in the human population in comparison to pandemic HIV-1 group M.

In summary, here we shed light on the evolutionary interplay between lentiviral Vif and host A3 in great apes and demonstrated that the interplay between Vif and A3 in GOR has uniquely affected the evolutionary episode of great ape lentiviruses. We elucidated how SIVcpz*Ptt* Vif overcame a species barrier mediated by GOR A3G. This is the first report explaining how great ape lentiviral Vif dismantled the hurdle mediated by new host A3 that potently impedes cross-species transmission of great ape lentiviruses, which has naturally occurred in Africa.

## STAR^*^METHODS

- KEY RESOURCES TABLE
- CONTACT FOR REAGENT AND RESOURCE SHARING
- EXPERIMENTAL MODEL AND SUBJECT DETAILS
  - Cells and Viruses
- METHOD DETAILS
  - Molecular Phylogenetic
  - Expression Plasmids
  - Western Blotting
  - TZM-bl Reporter Assay for Virus Infectivity Quantification
  - Multi-Round Virus Infection
- QUANTIFICATION AND STATISTICAL ANALYSIS

## Supplemental Information

Supplemental Information includes three figures five tables and can be found with this article online.

## Author Contributions

Y.N., K.Y., A.S., R.K., H.A., N.M., Y.Konno, I.K., S.N. and G.J.-F. performed the experiments and analyzed the data. J.I. and K.S. conducted phylogenetic analyses. T.I., Y.Koyanagi and R.S.H. provided reagents. K.S. conceived and designed the experiments. T.I., R.S.H. and K.S. wrote the manuscript.

## Acknowledgments

We would like to thank Beatrice H. Hahn and Frederic Bibollet-Ruche (University of Pennsylvania, USA) and Frank Kirchhoff and Daniel Sauter (Ulm University, Germany) for kindly providing the IMCs of some HIV-1, SIVcpz*Ptt* and SIVgor, and James L. Riley (University of Pennsylvania, USA) for sharing the CCR5-expressing lentivirus vector. We also thank Kotubu Misawa for dedicated support.

This study was supported in part by AMED J-PRIDE 18fm0208006h0002 (to K.S.), AMED Research on HIV/AIDS 18fk0410019h0001 (to K.S.) and 18fk0410014h0001 (to Y.Koyanagi), JST CREST (to K.S.), JSPS KAKENHI for Scientific Research B (Generative Research Fields) 16KT0111 (to K.S.), Scientific Research B 18H02662 (to K.S.) and Scientific Research on Innovative Areas 16H06429 (to K.S), 16K21723 (to K.S.) and 17H05813 (to K.S.), Takeda Science Foundation (to K.S.), Smoking Research Foundation (to K.S.), Mishima Kaiun Memorial Foundation (to K.S.), Tobemaki Foundation (to K.S.), ONO Medical Research Foundation (to K.S.), Joint Usage/Research Center program of Institute for Frontier Life and Medical Sciences, Kyoto University (to K.S.), and JSPS Core-to-Core program, A. Advanced Research Networks (to Y.Koyanagi, R.S.H. and K.S.), and NIAID R37 AI064046 (to R.S.H.). R.S.H. is the Margaret Harvey Schering Land Grant Chair for Cancer Research, a Distinguished McKnight University Professor, and an Investigator of the Howard Hughes Medical Institute.

## STAR*METHODS

### CONTACT FOR REAGENT AND RESOURSE SHARING

Further information and requests for resources and reagents may be directed to and will be fulfilled by the Lead Contact, Kei Sato.

### EXPERIMENTAL MODEL AND SUBJECT DETAILS

#### Cells and Viruses

HEK293T cells (a human embryonic kidney cell line; ATCC CRL-1573) and TZM-bl cells (obtained through the NIH AIDS Research and Reference Reagent Program) (Wei et al., 2002) were maintained in Dulbecco’s modified Eagle’s medium (Sigma) containing FCS and antibiotics. A T cell line SupT11 was prepared as reported previously (Refsland et al., 2010). To create the SupT11 cells stably expressing CCR5 (SupT11-CCR5), the CCR5-expressing lentivirus vector (Parry et al., 2003) and used for transduction. A single cell subclone of SupT11-CCR5 was isolated by limiting dilution. Surface expression levels of CD4, CCR5 and CXCR4 on SupT11-CCR5 cells were analyzed by staining with anti-CD4-FITC (Miltenyi Biotech; clone M-T466) or CD4-PE (Miltenyi Biotech; clone M-T466), CCR5-FITC (BD Pharmingen; clone 2D7) and CXCR4-PE (BD Pharmingen; clone 12G5) antibodies. To prepare the SupT11-CCR5 cells stably expressing GOR A3G (SupT11-CCR5-GOR A3G), a GOR A3G expression plasmid was transfected into SupT11-CCR5 cells using the Neon Transfection system (Thermo Fisher Scientific). The transfected cells were selected with Puromycin (InvivoGen; 0.5 ng/ml) and the expression of GOR A3G in the selected cell clone was analyzed by Western blotting using anti-HU A3G antibody (**Figure S2C**). Parental SupT11-CCR5 and SupT11-CCR5-GOR A3G cells were maintained in RPMI1640 (Sigma) containing FCS and antibiotics.

HEK293T cells were transfected using PEI Max (Polysciences) according to the manufacturer’s protocol. Basically, the expression plasmids for flag-tagged great ape A3D (HU, 50 ng; GOR, 200 ng), A3F (HU, 50 ng; GOR, 200 ng) or A3G (HU, 10 ng; GOR, 50 ng) were co-transfected with pNL4-3Δ*vif* (500 ng), an infectious molecular clone (IMC) of *vif*-deleted HIV-1M strain NL4-3 (Sato et al., 2014), and HA-tagged Vif expression plasmid (500 ng) into HEK293T cells. At two days post-transfection, the culture supernatants and transfected cells were harvested and were respectively used for TZM-bl assay and Western blotting as described below. For the preparation of infectious viruses, the IMCs (1,000 ng) were transfected into HEK293T cells. At two days post-transfection, the culture supernatants were harvested, centrifuged, and then filtered through a 0.45-μm-pore-size filter. To titrate virus infectivity, TZM-bl assay was performed as described below.

### METHOD DETAILS

#### Molecular Phylogenetic

The *vif* open reading frame (ORF) sequences (listed in **Table S1**) were extracted from Los Alamos National Laboratory HIV sequence database (https://www.hiv.lanl.gov/components/sequence/HIV/search/search.html) and aligned by using Clustal W implemented in MEGA 7 software (Kumar et al., 2016). Maximum-likelihood phylogenetic trees (Figures 1A, **S1A and S3A**) were constructed using MEGA 7 software (Kumar et al., 2016). The logoplot of Vif amino acid sequence is constructed using WebLogo 3 (http://weblogo.threeplusone.com) and the residues at positions 14-17 (DRMR motif) and 40-44 (YRHHY motif) are shown in Figure 4A.

#### Expression Plasmids

To construct the expression plasmids for flag-tagged great ape A3s, pcDNA3.1 plasmid (Thermo Fisher Scientific) was used as a backbone. The expression plasmids for flag-tagged HU A3D, A3F and A3G were used in our previous study (Nakano et al., 2017b). To construct the expression plasmids for flag-tagged CPZ and GOR A3s, the ORFs of CPZ A3D (JN247642), CPZ A3F (XM_525658), CPZ A3G (NM_001009001), GOR A3D (JN247649), GOR A3F (JN247640) and GOR A3G (AY639868) were synthesized by the GeneArt gene synthesis service (Thermo Fisher Scientific). The obtained DNA fragments were inserted into EcoRV-Not1 (for A3D and A3F) or EcoRI-Xho1 (for A3G) sites of pcDNA3.1 plasmid (Thermo Fisher Scientific). To construct the expression plasmids for HA-tagged lentiviral Vifs, pDON-AI plasmid (Takara) was used as a backbone. The expression plasmid for the HA-tagged Vifs of HIV-1M NL4-3 (M19921), SIVcpz*Ptt* MB897 (JN835461) and SIVcpz*Ptt* MT145 (JN835462) were used in our previous study (Sato et al., 2018).

The HA-tagged Vif ORFs of HIV-1M JRCSF (M38429), HIV-1N DJO0131 (AY532635), HIV-1O BCF183, HIV-1O RBF206, HIV-1P RBF168 and SIVgor CP2139 (FJ424866) were prepared by PCR using their IMCs as the templates and primers listed in **Table S2**. The HA-tagged Vif ORFs of HIV-1P U14788 (HQ179987), SIVcpz*Ptt* CAM3 (AF115393), SIVcpz*Ptt* DP943 (EF535993), SIVcpz*Ptt* LB7 (DQ373064), SIVgor CP684 (FJ424871) and SIVgor BPID2 (KP004994) were synthesized by the GeneArt gene synthesis service (Thermo Fisher Scientific). The obtained DNA fragments were inserted into BamHI-SalI site of pDON-AI plasmid (Takara). The Vif mutants of SIVcpz*Ptt* MB897 and other strains were prepared by the GeneArt site-directed mutagenesis system (Thermo Fisher Scientific) using the primers listed in **Table S3**. The expression plasmid for HA-tagged JRCSF Vif derivatives (Figure 4D) were prepared by PCR using their IMCs as the templates and primers listed in **Table S4**. The IMCs of SIVcpz*Ptt* MB897 derivatives, M16E and E2X (vif-deleted) were generated by mutagenesis/overlap extension PCR using the primers listed in **Table S5**. Nucleotide sequences were determined by a DNA sequencing service (Fasmac), and the sequence data were analyzed by Sequencher v5.1 software (Gene Codes Corporation).

#### Western Blotting

Western blotting was performed as previously described (Nakano et al., 2017b; Yamada et al., 2018) using the following antibodies: anti-HA (clone 3F10; Roche), anti-Flag (clone M2; Sigma-Aldrich), anti-p24 goat antiserum (ViroStat), anti-alpha-tubulin (TUBA; clone DM1A; Sigma-Aldrich); and anti-HU A3G rabbit antiserum (NIH AIDS Reagent Program catalog number #10201). For Western blotting of viral particles, 370 μl of viral supernatant was ultracentrifuged at 100,000 × g for 1 h at 4°C using a TL-100 instrument (Beckman), and the pellet was lysed with 1 × SDS buffer. Transfected cells were lysed with RIPA buffer (25 mM HEPES [pH 7.4], 50 mM NaCl, 1 mM MgCl_2_, 50 μM ZnCl_2_, 10% glycerol, 1% Triton X-100) containing a protease inhibitor cocktail (Roche).

#### TZM-bl Reporter Assay for Virus Infectivity Quantification

TZM-bl assay was performed as previously described (Nakano et al., 2017b; Yamada et al., 2018). Briefly, 10 μl of virus supernatant was inoculated into TZM-bl cells in 96-well plate (Nunc), and the β-galactosidase activity was measured by using the Galacto-Star mammalian reporter gene assay system (Thermo Fisher Scientific) and a 2030 ARVO X multi-label counter instrument (PerkinElmer) according to the manufacturers’ procedure. The relative infectivity was determined by relative light unit of this assay.

#### Multi-Round Virus Infection

The virus supernatant of SIVcpz*Ptt* MB897 WT, M16E and E2X (*vif*-deleted) derivatives were inoculated into SupT11-CCR5 (parental) and SupT11-CCR5-GOR A3G cells at multiplicity of infection (MOI) 0.1. The culture supernatant was harvested every two days and the amount of infectious viruses was measured by TZM-bl assay.

### QUANTIFICATION AND STATISTICAL ANALYSIS

Data analyses were performed using GraphPad Prism software. The data are presented as averages ± SEM. Statistically significant differences were determined by Student’s *t* test. Statistical details can be found directly in the figures or in the corresponding figure legends.

### STAR*METHODS

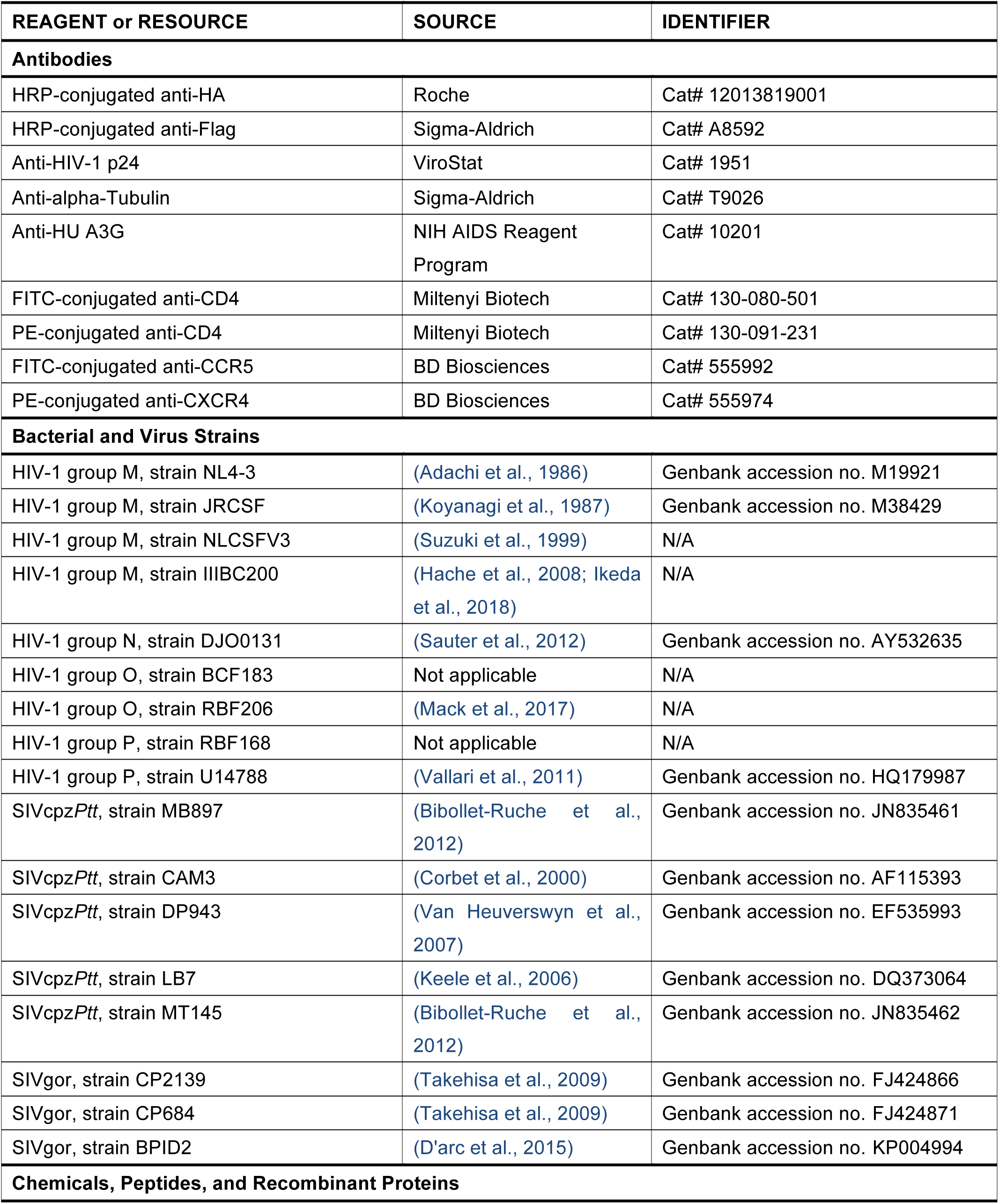
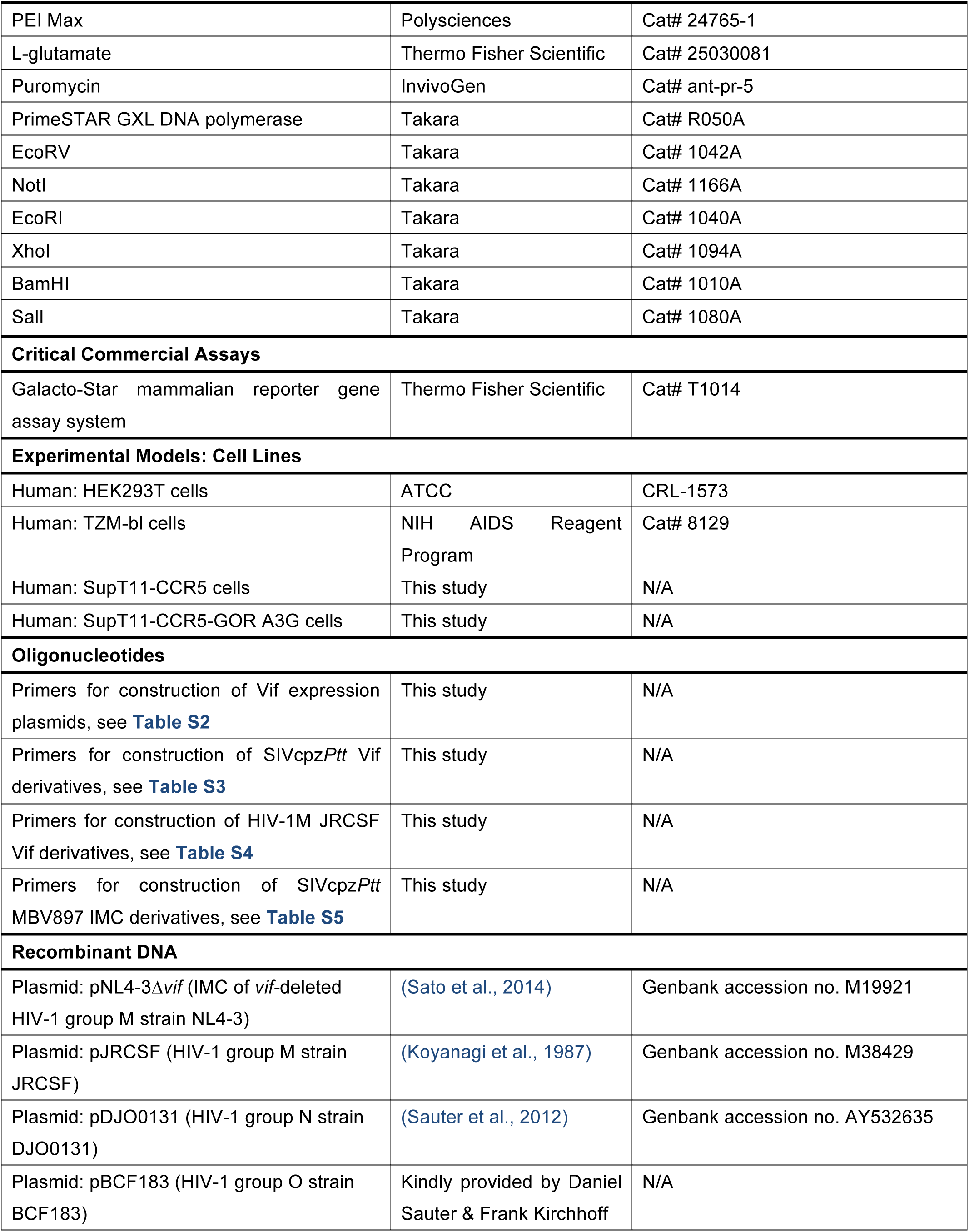
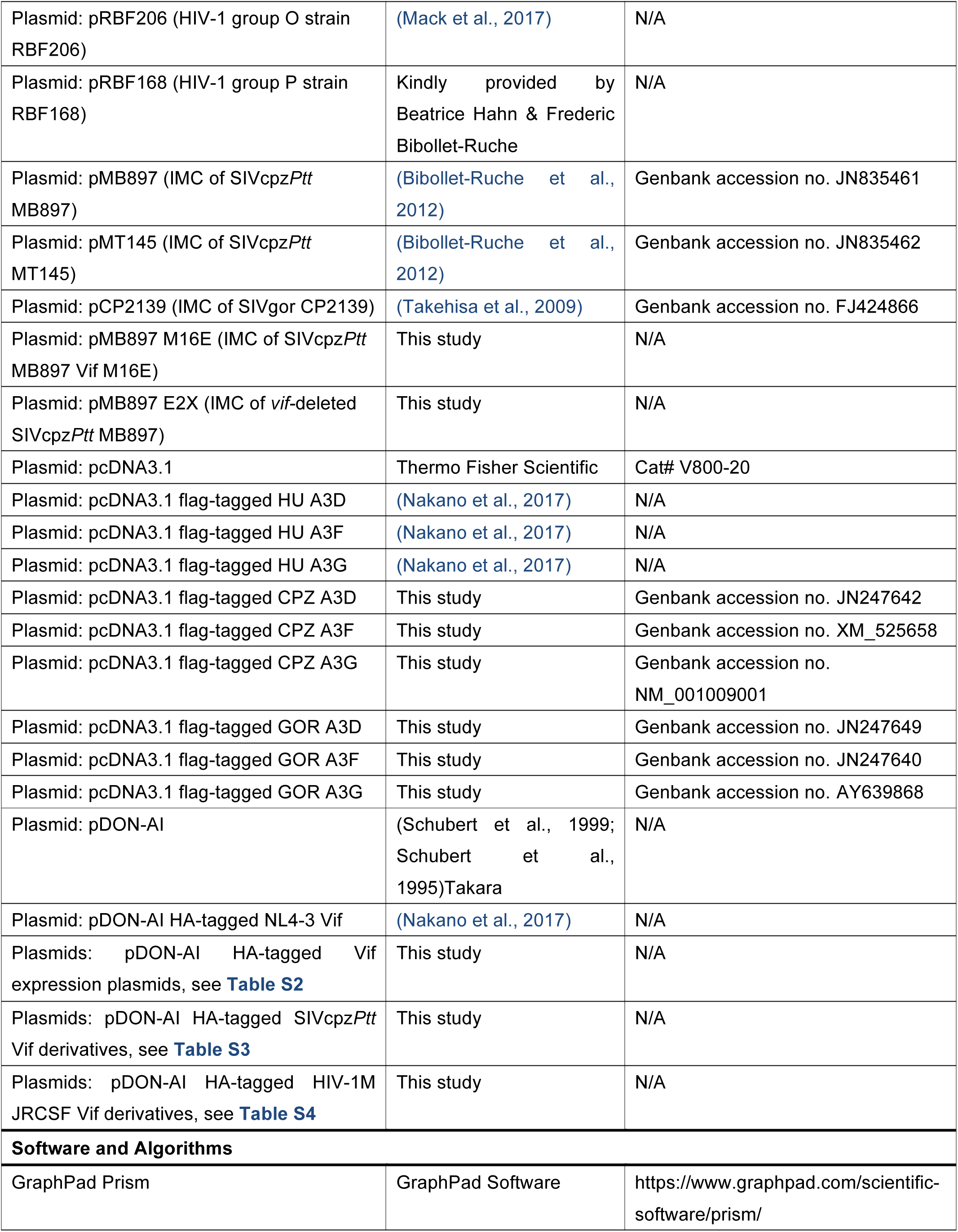
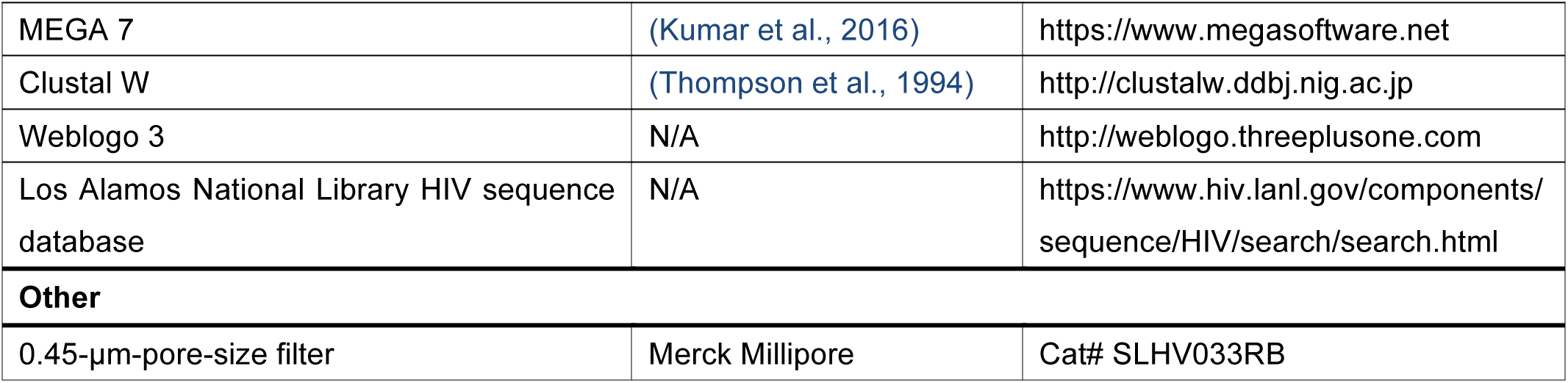
**KEY RESOURCES TABLE**

## References

Barre-Sinoussi, F., Chermann, J.C., Rey, F., Nugeyre, M.T., Chamaret, S., Gruest, J., Dauguet, C., Axler-Blin, C., Vezinet-Brun, F., Rouzioux, C., et al. (1983). Isolation of a T-lymphotropic retrovirus from a patient at risk for acquired immune deficiency syndrome (AIDS). Science 220, 868–871.

Bibollet-Ruche, F., Heigele, A., Keele, B.F., Easlick, J.L., Decker, J.M., Takehisa, J., Learn, G., Sharp, P.M., Hahn, B.H., and Kirchhoff, F. (2012). Efficient SIVcpz replication in human lymphoid tissue requires viral matrix protein adaptation. J Clin Invest 122, 1644–1652.

Compton, A.A., Bruel, T., Porrot, F., Mallet, A., Sachse, M., Euvrard, M., Liang, C., Casartelli, N., and Schwartz, O. (2014). IFITM proteins incorporated into HIV-1 virions impair viral fusion and spread. Cell Host Microbe 16, 736–747.

Compton, A.A., and Emerman, M. (2013). Convergence and divergence in the evolution of the APOBEC3G-Vif interaction reveal ancient origins of simian immunodeficiency viruses. PLoS Pathog 9, e1003135.

D’arc, M., Ayouba, A., Esteban, A., Learn, G.H., Boue, V., Liegeois, F., Etienne, L., Tagg, N., Leendertz, F.H., Boesch, C., et al. (2015). Origin of the HIV-1 group O epidemic in western lowland gorillas. Proc Natl Acad Sci U S A 112, E1343–1352.

Doyle, T., Goujon, C., and Malim, M.H. (2015). HIV-1 and interferons: who’s interfering with whom? Nat Rev Microbiol 13, 403–413.

Gao, F., Bailes, E., Robertson, D.L., Chen, Y., Rodenburg, C.M., Michael, S.F., Cummins, L.B., Arthur, L.O., Peeters, M., Shaw, G.M., et al. (1999). Origin of HIV-1 in the chimpanzee Pan troglodytes troglodytes. Nature 397, 436–441.

Harris, R.S., and Dudley, J.P. (2015). APOBECs and virus restriction. Virology 479-480, 131–145.

Keele, B.F., Van Heuverswyn, F., Li, Y., Bailes, E., Takehisa, J., Santiago, M.L., Bibollet-Ruche, F., Chen, Y., Wain, L.V., Liegeois, F., et al. (2006). Chimpanzee reservoirs of pandemic and nonpandemic HIV-1. Science 313, 523–526.

Kirchhoff, F. (2010). Immune evasion and counteraction of restriction factors by HIV-1 and other primate lentiviruses. Cell Host Microbe 8, 55–67.

Kluge, S.F., Mack, K., Iyer, S.S., Pujol, F.M., Heigele, A., Learn, G.H., Usmani, S.M., Sauter, D., Joas, S., Hotter, D., et al. (2014). Nef proteins of epidemic HIV-1 group O strains antagonize human tetherin. Cell Host Microbe 16, 639–650.

Kluge, S.F., Sauter, D., and Kirchhoff, F. (2015). SnapShot: antiviral restriction factors. Cell 163, 774–774 e771.

Kumar, S., Stecher, G., and Tamura, K. (2016). MEGA 7: Molecular Evolutionary Genetics Analysis Version 7.0 for Bigger Datasets. Mol Biol Evol 33, 1870–1874.

Letko, M., Booiman, T., Kootstra, N., Simon, V., and Ooms, M. (2015). Identification of the HIV-1 Vif and Human APOBEC3G Protein Interface. Cell Rep 13, 1789–1799.

Letko, M., Silvestri, G., Hahn, B.H., Bibollet-Ruche, F., Gokcumen, O., Simon, V., and Ooms, M. (2013). Vif proteins from diverse primate lentiviral lineages use the same binding site in APOBEC3G. J Virol 87, 11861–11871.

Nakano, Y., Aso, H., Soper, A., Yamada, E., Moriwaki, M., Juarez-Fernandez, G., Koyanagi, Y., and Sato, K. (2017a). A conflict of interest: the evolutionary arms race between mammalian APOBEC3 and lentiviral Vif. Retrovirology 14, 31.

Nakano, Y., Misawa, N., Juarez-Fernandez, G., Moriwaki, M., Nakaoka, S., Funo, T., Yamada, E., Soper, A., Yoshikawa, R., Ebrahimi, D., et al. (2017b). HIV-1 competition experiments in humanized mice show that APOBEC3H imposes selective pressure and promotes virus adaptation. PLoS Pathog 13, e1006348.

Neil, S.J., Zang, T., and Bieniasz, P.D. (2008). Tetherin inhibits retrovirus release and is antagonized by HIV-1 Vpu. Nature 451, 425–430.

Parry, R.V., Rumbley, C.A., Vandenberghe, L.H., June, C.H., and Riley, J.L. (2003). CD28 and inducible costimulatory protein Src homology 2 binding domains show distinct regulation of phosphatidylinositol 3-kinase, Bcl-xL, and IL-2 expression in primary human CD4 T lymphocytes. J Immunol 171, 166–174.

Refsland, E.W., Stenglein, M.D., Shindo, K., Albin, J.S., Brown, W.L., and Harris, R.S. (2010). Quantitative profiling of the full APOBEC3 mRNA repertoire in lymphocytes and tissues: implications for HIV-1 restriction. Nucleic Acids Res 38, 4274–4284.

Russell, R.A., Smith, J., Barr, R., Bhattacharyya, D., and Pathak, V.K. (2009). Distinct domains within APOBEC3G and APOBEC3F interact with separate regions of human immunodeficiency virus type 1 Vif. J Virol 83, 1992–2003.

Sato, K., Misawa, N., Takeuchi, J.S., Kobayashi, T., Izumi, T., Aso, H., Nagaoka, S., Yamamoto, K., Kimura, I., Konno, Y., et al. (2018). Experimental adaptive evolution of simian immunodeficiency virus SIVcpz to pandemic human immunodeficiency virus type 1 by using a humanized mouse model. J Virol 92, e01905.

Sato, K., Takeuchi, J.S., Misawa, N., Izumi, T., Kobayashi, T., Kimura, Y., Iwami, S., Takaori-Kondo, A., Hu, W.S., Aihara, K., et al. (2014). APOBEC3D and APOBEC3F potently promote HIV-1 diversification and evolution in humanized mouse model. PLoS Pathog 10, e1004453.

Sauter, D., Schindler, M., Specht, A., Landford, W.N., Munch, J., Kim, K.A., Votteler, J., Schubert, U., Bibollet-Ruche, F., Keele, B.F., et al. (2009). Tetherin-driven adaptation of Vpu and Nef function and the evolution of pandemic and nonpandemic HIV-1 strains. Cell Host Microbe 6, 409–421.

Sharp, P.M., and Hahn, B.H. (2011). Origins of HIV and the AIDS pandemic. Cold Spring Harb Perspect Med 1, a006841.

Takehisa, J., Kraus, M.H., Ayouba, A., Bailes, E., Van Heuverswyn, F., Decker, J.M., Li, Y., Rudicell, R.S., Learn, G.H., Neel, C., et al. (2009). Origin and biology of simian immunodeficiency virus in wild-living western gorillas. J Virol 83, 1635–1648.

Van Damme, N., Goff, D., Katsura, C., Jorgenson, R.L., Mitchell, R., Johnson, M.C., Stephens, E.B., and Guatelli, J. (2008). The interferon-induced protein BST-2 restricts HIV-1 release and is downregulated from the cell surface by the viral Vpu protein. Cell Host Microbe 3, 245–252.

Van Heuverswyn, F., Li, Y., Neel, C., Bailes, E., Keele, B.F., Liu, W., Loul, S., Butel, C., Liegeois, F., Bienvenue, Y., et al. (2006). Human immunodeficiency viruses: SIV infection in wild gorillas. Nature 444, 164.

Vanden Haesevelde, M.M., Peeters, M., Jannes, G., Janssens, W., van der Groen, G., Sharp, P.M., and Saman, E. (1996). Sequence analysis of a highly divergent HIV-1-related lentivirus isolated from a wild captured chimpanzee. Virology 221, 346–350.

Wei, X., Decker, J.M., Liu, H., Zhang, Z., Arani, R.B., Kilby, J.M., Saag, M.S., Wu, X., Shaw, G.M., and Kappes, J.C. (2002). Emergence of resistant human immunodeficiency virus type 1 in patients receiving fusion inhibitor (T-20) monotherapy. Antimicrob Agents Chemother 46, 1896–1905.

Yamada, E., Nakaoka, S., Klein, L., Reith, E., Langer, S., Hopfensperger, K., Iwami, S., Schreiber, G., Kirchhoff, F., Koyanagi, Y., et al. (2018). Human-Specific Adaptations in Vpu Conferring Anti-tetherin Activity Are Critical for Efficient Early HIV-1 Replication In Vivo. Cell Host Microbe 23, 110–120 e117.

Zhang, Z., Gu, Q., de Manuel Montero, M., Bravo, I.G., Marques-Bonet, T., Haussinger, D., and Munk, C. (2017). Stably expressed APOBEC3H forms a barrier for cross-species transmission of simian immunodeficiency virus of chimpanzee to humans. PLoS Pathog 13, e1006746.

## References

Adachi, A., Gendelman, H.E., Koenig, S., Folks, T., Willey, R., Rabson, A., and Martin, M.A. (1986). Production of acquired immunodeficiency syndrome-associated retrovirus in human and nonhuman cells transfected with an infectious molecular clone. J Virol 59, 284–291.

Corbet, S., Muller-Trutwin, M.C., Versmisse, P., Delarue, S., Ayouba, A., Lewis, J., Brunak, S., Martin, P., Brun-Vezinet, F., Simon, F., et al. (2000). env sequences of simian immunodeficiency viruses from chimpanzees in Cameroon are strongly related to those of human immunodeficiency virus group N from the same geographic area. J Virol 74, 529–534.

Hache, G., Shindo, K., Albin, J.S., and Harris, R.S. (2008). Evolution of HIV-1 isolates that use a novel Vif-independent mechanism to resist restriction by human APOBEC3G. Curr Biol 18, 819–824.

Ikeda, T., Symeonides, M., Albin, J.S., Li, M., Thali, M., and Harris, R.S. (2018). HIV-1 adaptation studies reveal a novel Env-mediated homeostasis mechanism for evading lethal hypermutation by APOBEC3G. PLoS Pathog 14, e1007010.

Koyanagi, Y., Miles, S., Mitsuyasu, R.T., Merrill, J.E., Vinters, H.V., and Chen, I.S. (1987). Dual infection of the central nervous system by AIDS viruses with distinct cellular tropisms. Science 236, 819–822.

Kumar, S., Stecher, G., and Tamura, K. (2016). MEGA7: Molecular Evolutionary Genetics Analysis Version 7.0 for Bigger Datasets. Mol Biol Evol 33, 1870–1874.

Mack, K., Starz, K., Sauter, D., Langer, S., Bibollet-Ruche, F., Learn, G.H., Sturzel, C.M., Leoz, M., Plantier, J.C., Geyer, M., et al. (2017). Efficient Vpu-Mediated Tetherin Antagonism by an HIV-1 Group O Strain. J Virol 91.

Nakano, Y., Misawa, N., Juarez-Fernandez, G., Moriwaki, M., Nakaoka, S., Funo, T., Yamada, E., Soper, A., Yoshikawa, R., Ebrahimi, D., et al. (2017). HIV-1 competition experiments in humanized mice show that APOBEC3H imposes selective pressure and promotes virus adaptation. PLoS Pathog 13, e1006348.

Sauter, D., Unterweger, D., Vogl, M., Usmani, S.M., Heigele, A., Kluge, S.F., Hermkes, E., Moll, M., Barker, E., Peeters, M., et al. (2012). Human tetherin exerts strong selection pressure on the HIV-1 group N Vpu protein. PLoS Pathog 8, e1003093.

Schubert, U., Bour, S., Willey, R.L., and Strebel, K. (1999). Regulation of virus release by the macrophage-tropic human immunodeficiency virus type 1 AD8 isolate is redundant and can be controlled by either Vpu or Env. J Virol 73, 887–896.

Schubert, U., Clouse, K.A., and Strebel, K. (1995). Augmentation of virus secretion by the human immunodeficiency virus type 1 Vpu protein is cell type independent and occurs in cultured human primary macrophages and lymphocytes. J Virol 69, 7699–7711.

Suzuki, Y., Koyanagi, Y., Tanaka, Y., Murakami, T., Misawa, N., Maeda, N., Kimura, T., Shida, H., Hoxie, J.A., O’Brien, W.A., et al. (1999). Determinant in human immunodeficiency virus type 1 for efficient replication under cytokine-induced CD4+ T-helper 1 (Th1)- and Th2-type conditions. J Virol 73, 316–324.

Thompson, J.D., Higgins, D.G., and Gibson, T.J. (1994). CLUSTAL W: improving the sensitivity of progressive multiple sequence alignment through sequence weighting, position-specific gap penalties and weight matrix choice. Nucleic Acids Res 22, 4673–4680.

Vallari, A., Holzmayer, V., Harris, B., Yamaguchi, J., Ngansop, C., Makamche, F., Mbanya, D., Kaptue, L., Ndembi, N., Gurtler, L., et al. (2011). Confirmation of putative HIV-1 group P in Cameroon. J Virol 85, 1403–1407.

Van Heuverswyn, F., Li, Y., Bailes, E., Neel, C., Lafay, B., Keele, B.F., Shaw, K.S., Takehisa, J., Kraus, M.H., Loul, S., et al. (2007). Genetic diversity and phylogeographic clustering of SIVcpz*Ptt* in wild chimpanzees in Cameroon. Virology 368, 155–171.

